# The unphosphorylated form of the PAQosome core subunit RPAP3 binds ribosomal preassembly complexes to modulate ribosome biogenesis

**DOI:** 10.1101/2021.03.17.435124

**Authors:** Maxime Pinard, Philippe Cloutier, Christian Poitras, Marie-Soleil Gauthier, Benoit Coulombe

## Abstract

The PAQosome (Particle for Arrangement of Quaternary structure) is a twelve-subunit HSP90 co-chaperone involved in the biogenesis of several human protein complexes. Two mechanisms of client selection have previously been identified, namely the selective recruitment of specific adaptors and the differential use of homologous core subunits. Here, we describe a third client selection mechanism by showing that RPAP3, one of the core PAQosome subunits, is phosphorylated at several Ser residues in HEK293 cells. Affinity purification coupled with mass spectrometry (AP-MS) using expression of tagged RPAP3 with single phospho-null mutations at Ser116, Ser119 or Ser121 reveals binding of the unphosphorylated form to several proteins involved in ribosome biogenesis. *In vitro* phosphorylation assays indicate that the kinase CK2 phosphorylates these RPAP3 residues. This finding is supported by data showing that pharmacological inhibition of CK2 enhances binding of RPAP3 to ribosome preassembly factors in AP-MS experiments. Moreover, silencing of PAQosome subunits interferes with ribosomal assembly factors’ interactome. Altogether, these results indicate that RPAP3 phosphate group addition/removal at specific residues modulates binding to subunits of preribosomal complexes and allows speculating that PAQosome posttranslational modifications is a mechanism of client selection.

## Introduction

The PAQosome (also known as R2TP/PFDL; *<u>R</u>vb1p, <u>R</u>vb2p, <u>T</u>ah1p and <u>P</u>ih1p/prefoldin-like*) is a 12-subunit chaperone complex responsible for the biogenesis of a vast number of protein complexes in humans^1, 2^. Its *Saccharomyces cerevisiae* counterpart, which is apparently lacking the prefoldin-like (PFDL) module, was first identified based on its association with Hsp90^3^ and then shown to orchestrate the assembly of snoRNPs, ribonucleoprotein (RNP) complexes involved in the maturation of ribosomal rRNA^4, 5^ while new cofactor participating in the snoRNP complex formation were recently identified^6^. This chaperone complex was also observed in association with subunits of all three nuclear RNA polymerases and likewise promotes their cytoplasmic assembly before translocation of mature complexes to different compartments of the nucleus^7, 8^. More recently, a role in the biogenesis of spliceosome component U4 and U5 snRNPs was revealed^9–11^. In addition to targeting protein and RNP complexes that mediate key steps of gene expression from transcription to translation, the PAQosome was shown to affect the cellular response to stress through stabilization of phosphatidylinositol 3-kinase-related kinases (PIKK)^12^. This stress response promotes the assembly of various complexes involved in nutrient sensing and growth factor signaling pathway (mTOR)^13, 14^, DNA damage response (DNA-PK, ATM and ATR)^12, 14, 15^, chromatin remodelling (TRRAP)^12^ and non-sense mediated mRNA decay (SMG-1)^13^.

Two mechanisms of client selection by the PAQosome have been identified so far. The first one consists of specific clients being recruited to the PAQosome, some directly^7, 8^ and some indirectly via adaptor proteins^4–6, 9–11, 13^, and can then be assembled and stabilized by the platform formed by the PAQosome together with HSP90. Moreover, it was shown that phosphorylation can affect the interaction between some adaptors and subunits of the R2TP/Prefoldin-like complex such as that of TEL2 with PIHD1^13^. As our cells must assemble thousands of protein complexes in a dynamically regulated manner, it is highly improbable that the original PAQosome is the only cellular particle responsible for assembly of all these complexes. Recently, Bertrand and colleagues discovered a second client selection mechanism by showing that some PAQosome subunits have homologous proteins that can form alternative PAQosome complexes with their own specific sets of clients^16^. Moreover, the data support the notion that alternative PAQosomes can harbor tissue specificity. Indeed, a version of the PAQosome in which RPAP3 is replaced by the homologous protein SPAG1, and PIH1D1 by PIH1D2, is enriched in testis^16^. These modest differences drastically alter the specificity of the resulting particle, which is now able to recognize a totally different set of clients such as liprin complexes^16^ or those involved in dynein arms formation^17^. This work on other PIH1D1 and RPAP3 isoforms and homologous proteins, together with data detailing the contribution of several adaptors, allows speculating that the PAQosome exists in many other flavors, each assembling a number of specific complexes and networks.

In the current study, we report on the identification of a third mechanism of substrate selection by the PAQosome. By exploring the function of PAQosome phosphorylation, we found that phosphorylation of the core subunit RPAP3 regulates client binding. Indeed, RPAP3 is found to bind ribosomal assembly factors when this core PAQosome subunit is in the unphosphorylated state. It will be interesting to assess in the future whether various posttranslational modifications of RPAP3 or other PAQosome subunits also participate in client selection.

## Experimental section

### DNA plasmid constructs

All tagged constructs used in AP-MS and GST pulldown experiments were produced by PCR amplification of PIH1D1, RPAP3, RRP1B and PDCD11 cDNA using Mammalian Gene Collection clones (GE Healthcare) using specific oligonucleotides (Supplementary Table S1). The resulting fragments were subsequently cloned into p3xFLAG-CMV-14 (Sigma) or pGEX-6P-1 (GE Healthcare) and transformed in One Shot OmniMAX 2 T1R Chemically Competent *E. coli* (ThermoFisher, C854003). Plasmid purification was done using QIAprep Spin Miniprep Kit (QIAGEN, 271060) and the sequences were verified by sequencing. Phospho-null and phosphomimetic mutants were introduced in RPAP3 by site-directed mutagenesis with specific oligonucleotides and plasmid sequences were validated by sequencing (Supplementary Table S1).

### FLAG affinity purification

This protocol was modified from Kean et al.^18^. PIH1D1-WT and RPAP3-WT or the various RPAP3 mutant (S116A, S119A and S121A) expression vectors were transiently transfected in two wells of a 6-well plate containing HEK 293T cells (ATCC, CRL-3216) at 60% confluency using jetPRIME (Polyplus, 114) according to manufacturer’s instructions. The empty vector was used as a negative control. Cells were incubated in complete DMEM containing 4,5g/L glucose (ThermoFisher, 11995-065) supplemented with 10% fetal bovine serum (Wisent, 080-150), 2 mM glutamine (ThermoFisher, 25030081), 100 U/mL penicillin and 100 µg/mL streptomycin (ThermoFisher, 15140122) and harvested 48 h later. PIH1D1-WT transfected cells were incubated with starved conditions (DMEM containing 1g/L glucose (ThermoFisher, 11885-084) supplemented with 2 mM glutamine, 100 U/mL penicillin and 100 µg/mL streptomycin). In indicated experiments, 10µM of CX-4945 (SelleckChem, S2248) or 100µM of TBB (Selleckchem, S5265) was added 32 h after RPAP3-WT transfection and cells were harvested at 48 h post transfection. DMSO-treated cells were used as negative controls. In the case of siRNA treated purifications, cells were grown to a confluence of 20% and transfected with SMARTpool siGenome siRNA targeting human RPAP3 (Dharmacon, M-014385-00-0005), human WDR92 (Dharmacon, M-008669-01-0005) or a non-targeting siRNA control (Dharmacon, D-001210-01) at a concentration of 100 nM. The following day, cells were cotransfected again with siRNAs and either the 3xFLAG-RRP1B or 3xFLAG-PDCD11 expression vector and harvested 48 h later. Cell membranes were disrupted by two liquid nitrogen freeze-thaw cycles in 1 mL lysis buffer (100 mM KCl, 50 mM HEPES-KOH pH 8.0, 2 mM EDTA, 0.1% NP-40, 10% glycerol, 1 mM DTT and cOmplete EDTA-free Protease Inhibitor Cocktail (MilliporeSigma, COEDTAF-RO)). The protein extracts were cleared of insoluble material by centrifugation (12,000 g, 30 min, 4 °C). 20µL of anti-FLAG M2 magnetic bead slurry (MilliporeSigma, M8823) were washed twice 1 mL of lysis buffer, then incubated for three hours with 1,5 to 2 mg of total proteins at 4 °C on a tube rotator. The beads were then washed three times in 1 mL lysis buffer and three more times in 1 mL wash buffer (75 mM KCl, 50 mM ammonium bicarbonate pH 8.0). The bound proteins were eluted by three successive 30 min incubations in 300 µl of ammonium hydroxide solution pH 11-12 (roughly 10%) in HPLC-grade water (Sigma, 7732-18-5). The combined fractions were dried in a speed-vac and then resuspended in 10 µL of 6M urea. Reduction buffer was added at a volume of 2.5 µL (45 mM DTT, 100 mM ammonium bicarbonate) and the samples were incubated for 30 min at 37 °C. An additional 2.5 µL of alkylation buffer (100 mM iodoacetamide, 100 mM ammonium bicarbonate) were included to the mix followed by 20 min incubation at 24 °C and in absence on light. Twenty microliters of HPLC-grade water were added to reduce urea concentration and trypsin digestion was performed using a 1:20 (enzyme:protein) ratio of sequencing grade modified trypsin (Promega, V5111) and 18 h incubation at 37 °C on a ThermoMixer (Eppendorf, 5382000023). Following a quick centrifugation (500 g, 1 min) the peptides were collected. For most samples, trifluoroacetic acid was added to the samples and residual salts and detergents were removed using an Oasis MCX 96-well μElution Plate (Waters, 186001830BA) loaded onto a Positive Pressure-96 Processor (Waters, 186006961) according to the manufacturer’s instructions. 3XFLAg-PIH1D1 samples were re-solubilized under agitation for 15 min in 10 µL of 0.2% formic acid. Desalting/cleanup of the digests was performed by using C18 ZipTip pipette tips (Millipore, Billerica, MA). Eluates were dried down in vacuum centrifuge and then re-solubilized under agitation for 15 min in 12 µL of 2% acetonitrile, 1% formic acid, of which 5 µL were used for LC-MS/MS.

### Mass spectrometry

High performance liquid chromatography (HPLC) was performed using C18 resin (5 µm particles, 300 Å pores) extracted from a Jupiter LC Column (00B-4053-E0) and packed in a PicoFrit Column (NEW OBJECTIVE, PF360-75-15-N-5), the latter of which was loaded into a Easy-nLC II instrument (ThermoFisher). Employed buffers were: 0.2% formic acid (buffer A) and 100% acetonitrile/0.2% formic acid (buffer B). Peptide elution was performed with a two-slope gradient at a flowrate of 250 nL/min. Solvent B gradually increased from 2 to 37% over the course of 90 min and then from 37 to 80% B in 10 min. The HPLC system was coupled to an Orbitrap Fusion Tribrid mass spectrometer (Thermo Scientific, IQLAAEGAAPFADBMBCX) through a nano-ESI source (ThermoFisher, ES071). Nanospray and S-lens voltages were set to 1.3–1.8 kV and 50 V, respectively. Capillary temperature was set to 225 °C. Full scan MS survey spectra (m/z 360-1560) in profile mode were acquired in the Orbitrap with a resolution of 120,000 with a target value at 1e6. The 25 most intense peptide ions were fragmented in the HCD collision cell and analyzed in the linear ion trap with a target value at 2e4 and normalized collision energy at 28. Target ions selected for fragmentation were dynamically excluded for 25 sec. The peak list files were generated with Proteome Discoverer v2.3 (ThermoFisher, OPTON-30812) using the following parameters: minimum mass set to 500 Da, maximum mass set to 6 kDa, no grouping of MS/MS spectra, precursor charge set to auto, and minimum number of fragment ions set to 5. Mass spectrometry experimental condition used for 3X-FLAG-PIH1D1-digested sample were previously described in Cloutier et al.^9^

### Data analysis

Label-Free Quantification (LFQ) and normalization was performed using MaxQuant v1.6.2.10^19^ and the characterized human UniProtKB database (release on June 3th 2018). Trypsin was chosen as digestion enzyme allowing a maximum of 2 miscleavages. Methionine oxidation and N-terminal acetylation where selected as variable modifications while cysteine carbamidomethylation was set as fixed modification. Fast LFQ was enabled and results were refined through the re-quantify option. Orbitrap was selected as instrument type with default parameters (First search peptide tolerance of 20 ppm; Main search tolerance of 4.5 ppm; Isotope match tolerance of 2 ppm; Centroid match tolerance of 8 ppm; Centroid half width of 35 ppm; Time valley factor: 1.4; isotope valley factor: 1.2; Isotope time correlation: 0.6; Theoretical isotope correlation: 0.6; Minimum peak length: 2; Maximum charge: 7; Minimum score for recalibration: 70; Gap scan of 1). Peptide identification was transferred across LC-MS runs based on similar m/z ratios and retention time using the “Match between runs” feature. Further statistical analysis was carried out with Perseus (version 1.6.1.3)^20^. All purifications were done in triplicate and proteins detected in two out of three experiments were kept for further analysis. Missing values were replaced by randomly generated intensities normally distributed with a width of 0.3 times and a downshift of 1.8 times the standard deviation of non-zero intensities. Significant differences between protein intensities from bait purifications and corresponding controls groups were then determined using a two-tailed T-test subsequently adjusted for multiple hypothesis testing with a permutation-based false discovery rate (FDR) of 0.05 and a fudge factor (s0) of 0.1 with 10,000 iterations. In the case of mutant comparaison with their WT counterpart (URI1 S372A/WT, RPAP3 S116A/WT, RPAP3 S119A/WT, RPAP3 S121A/WT) or drug-treated samples purifications, proteins that did not show enrichment in either conditions compared to the empty vector control (i.e. p-values over 0.05 and/or protein intensity ratio under 2) were discarded. Differential co-purification of the resulting subset of proteins in either condition was then carried out as detailed for bait versus control experiments. FLAg-tagged PIH1D1 purified protein search was performed with Mascot 2.5 (Matrix Science) against the Refseq Human protein database (Txid 9606) and against the R2TP sequence. The peak list files were generated with Proteome Discoverer (version 2.3) using the following parameters: minimum mass set to 500 Da, maximum mass set to 6000 Da, no grouping of MS/MS spectra, precursor charge set to auto, and minimum number of fragment ions set to 5. The mass tolerances for precursor and fragment ions were set to 10 ppm and 0.6 Da, respectively. Trypsin was used as the enzyme allowing for up to 2 missed cleavage. Cysteine carbamidomethylation was specified as a fixed modification, and methionine oxidation, serine, threonine and tyrosine phosphorylation modifications as variable modifications. Data analysis was performed using Scaffold (version 4.7) to select peptide containing phosphorylated residue. Spectrum was retrieve by using Tune software (Version 2.0). Gene ontology (GO) resource (http://geneontology.org/) using Panther software (version14.1)^21^ was used to annotate the proteins and perform a pathway analysis between RPAP3 Wt and one of the different phosphosite mutant (S116A, S119A or S121A). The proteins were annotated for the biological process. Benjamini-Hochberg false discovery rate (FDR) was obtain for all the significantly enrich protein obtain from the Log2 difference between RPAP3 WT and the different phosphosite mutant. The GO analysis was done against the GO database released 2020-03-23.

### GST-tagged protein purification

RPAP3-WT or various mutant (S116A, S119A and S121A and AAA) expressing vector transformation were performed with The resulting expression vector was used in the transformation of One Shot BL21 Star DE3 chemically competent *E. coli* (ThermoFisher, C601003) according to manufacturer’s specifications. Bacteria were then grown in 500 mL of 2xYT medium (16 g/L tryptone, 10 g/L yeast extract, 5.0 g/L NaCl, 100 µg/mL ampicillin) at 37 °C until culture density corresponding to an absorbance of 0.6 at 600 nm (OD600) was reached. Recombinant protein production was induced through addition of IPTG at a final concentration of 0.5 mM. After two hours of incubation, bacteria were harvested by centrifugation (3,000 g, 30 min, 4 °C) and lysed in 25 mL of PBS supplemented with 10 mg/mL lysozyme and cOmplete EDTA-free Protease Inhibitor Cocktail (MilliporeSigma, COEDTAF-RO) by running twice into a cooled French Press cell disruptor (ThermoFisher, FA-078A and FA-032). Lysates were cleared of insoluble material by ultracentrifugation (150,000g, 45 min, 4 °C) and the supernatant was transferred to a 50 mL tube containing 250 µl of Glutathione Sepharose 4B beads (GE Healthcare, 17075601) washed twice beforehand in 50 mL of PBS. The beads were incubated for 2 h at 4 °C on a tube rotator, washed three times with 50 mL of PBS and then transferred to 1.5 mL tubes. Proteins were then recovered with sequential 30 min incubations with 500 µL of elution buffer (10 mM Glutathione, 50 mM Tris–HCl pH 8.0). Elution efficiency was assessed by running fractions on SDS-PAGE followed by Coomassie staining. Fractions showing the strongest amounts of recombinant proteins were dialyzed overnight at 4 °C in storage and concentration buffer (100 mM KCl, 20 mM HEPES pH 7.9, 0.2 mM EDTA, 0.2 mM EGTA, 20% glycerol, 20% polyethylene glycol 8000 and 2 mM DTT). GST tagged was removed by using PreScission protease (sigma, GE27-0843-01) according to manufacturer’s specifications.

### *In vitro* phosphorylation assay

2µg of GST-cleaved RPAP3-WT or mutant (S116A, S119A, S121A or triple mutant (AAA)) was incubated and labeled *in vitro* with 2U of CK2 (New England Biolabs, P60105) in a 30µl reaction containing 3pM adenosine triphosphate (ATP), 50 mM Tris-HCl pH 7.5, 10 mM MgCl2, 0,1mM EDTA, 2mM DTT, 0,01% Brij35 and 5µCi[γ-32P]ATP for 30 min at 30°C. Reactions were stopped with sodium dodecyl sulfate polyacrylamide gel electrophoresis (SDS-PAGE) sample buffer. The samples were boiled and subjected to electrophoresis on a 10% SDS-PAGE. The stained SDS-PAGE gel was dried and the phosphorylation profile was obtained by autoradiography.

## Results

### Changes in the RPAP3 phosphorylation status regulate binding to preribosomal complexes

Posttranslational modification of PAQosome subunits stands out as a possible mechanism of client selection by this co-chaperone complex. Another group has shown that interaction of the PAQosome subunit URI1 with phosphatase 1 is increased upon URI1 dephosphorylation^22^. To address our hypothesis, we assess other PAQosome subunits for potential phosphorylation sites. LC-MS/MS analysis of affinity purified RPAP3 revealed phosphorylated Ser residues (S116 and S119) that are clustered near a sequence with homology to a putative nucleolus localization signal (NoLS) (Fig. 1A, 1B). The PTM database PhosphoSitePlus (https://www.phosphosite.org/homeAction.action) reports on the identification of two additional phosphorylation sites in this region, S87 and S121. Examination of the amino acid sequence surrounding these sites indicates that the sequence surrounding RPAP3 S87 has significant homology with the residues preceding URI1 S372 which corresponds to mTOR effector kinase S6K1 consensus recognition site R/K-x-R/K-x-x-S/T^22^. However, RPAP3 S119 and S121 bear more similarity to the Casein Kinase 2 (CK2) consensus site S-x-x-E/D^24^ (Fig. 1C). While the latter does not fit this pattern perfectly, prior phosphorylation of S119 may promote recognition of S116 by CK2, a situation reminiscent to that describes by the hierarchical model of St-Denis et al.^24^

**Fig 1.**
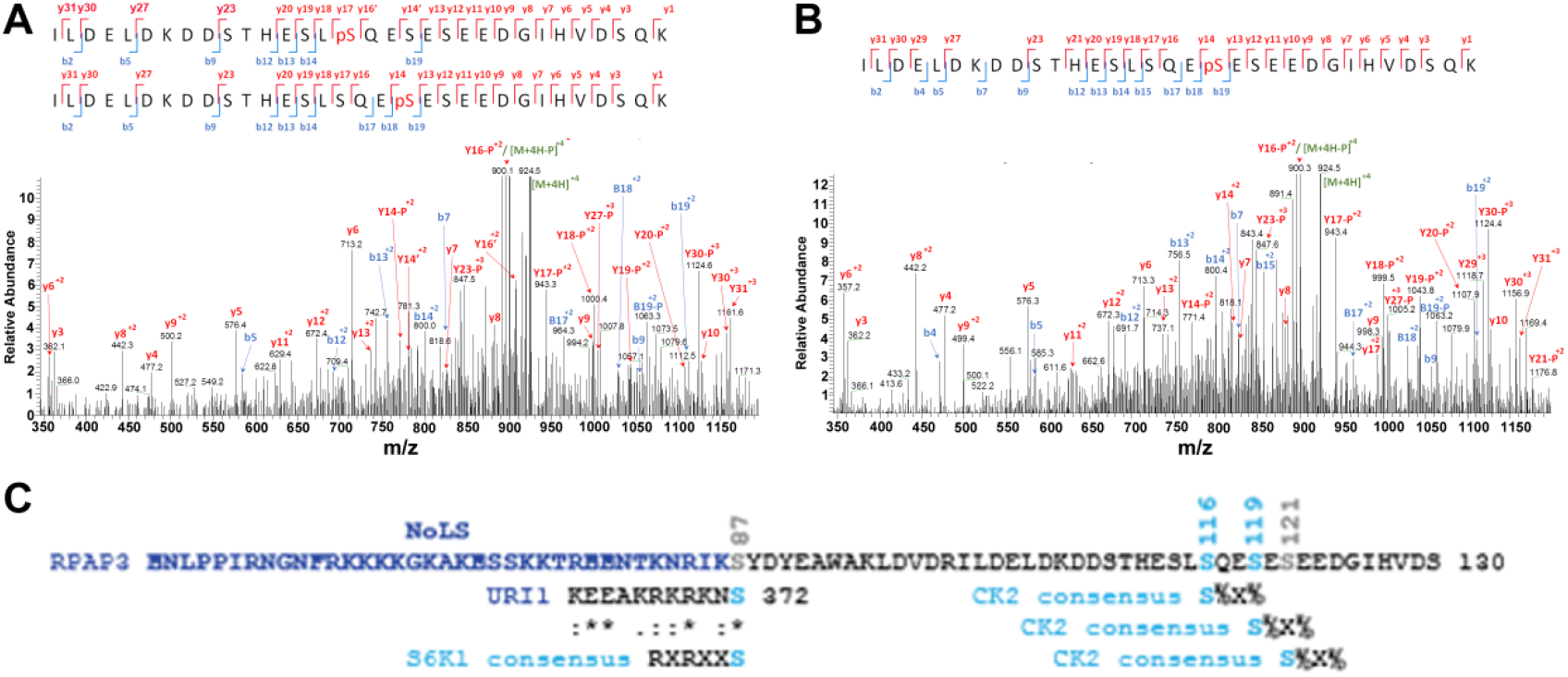
RPAP3 is phosphorylated at serine 116, 119 and 121, in close proximity to a putative nucleolar localization signal (NoLS). A) Annotated HCD MS/MS spectrum of RPAP3 peptide ILDELDKDDS THESLSQESES EEDGIHVDSQK monophosphorylated on S116 or S119. The precursor m/z value is 924.39953 (M+4H)+4, mass accuracy is −0.5 ppm and Mascot score is 33. B) Annotated HCD MS/MS spectrum of same peptide monophosphorylated at position S119. Precursor m/z value is 924.39842 (M+4H), mass accuracy is −1.3 ppm and Mascot score is 53. C) Sequence of the corresponding region of RPAP3 where monophosphorylated residues were observed (light blue; position is indicated above) as well as S87 and S121, two other phosphosites reported on PTM database PhosphoSitePlus with >30 entries^23^. The sequence surrounding RPAP3 S87 shows significant homology with the residues preceding URI1 S372 and corresponds to mTOR effector kinase S6K1 consensus recognition site, which has already been shown to be responsible for URI1 S372 phosphorylation^22^. RPAP3 S119 and S121, on the other hand, bear more similarity to Casein Kinase 2 (CK2) consensus site^24^.

To start elucidating the function of these phosphorylation sites, we use both phospho-null and phosphomimetic mutants in AP-MS experiments. To our surprise, phospho-null mutations (alanine substitution) of RPAP3 S116, S119 and S121 significantly increased the association of RPAP3 with several preribosomal structural proteins as well as proteins associated with ribosomal maturation^25^ (Fig. 2A, 2C, 2E; Supplementary Table S2, S3, S4). Phosphomimetic mutants of RPAP3 S116, S119 and S121 (Asp substitution) *versus* wild type RPAP3 did not result in a difference as marked as what phospho-null mutations showed in favor of preribosomal proteins (Fig. 2B, 2D, 2F; Supplementary Table S2, S3, S4). All mutants showed significant association with ribosomal proteins and biogenesis factors associated with small (40S; light blue) and large (60S; dark blue) subunits of the ribosome.

**Fig 2.**
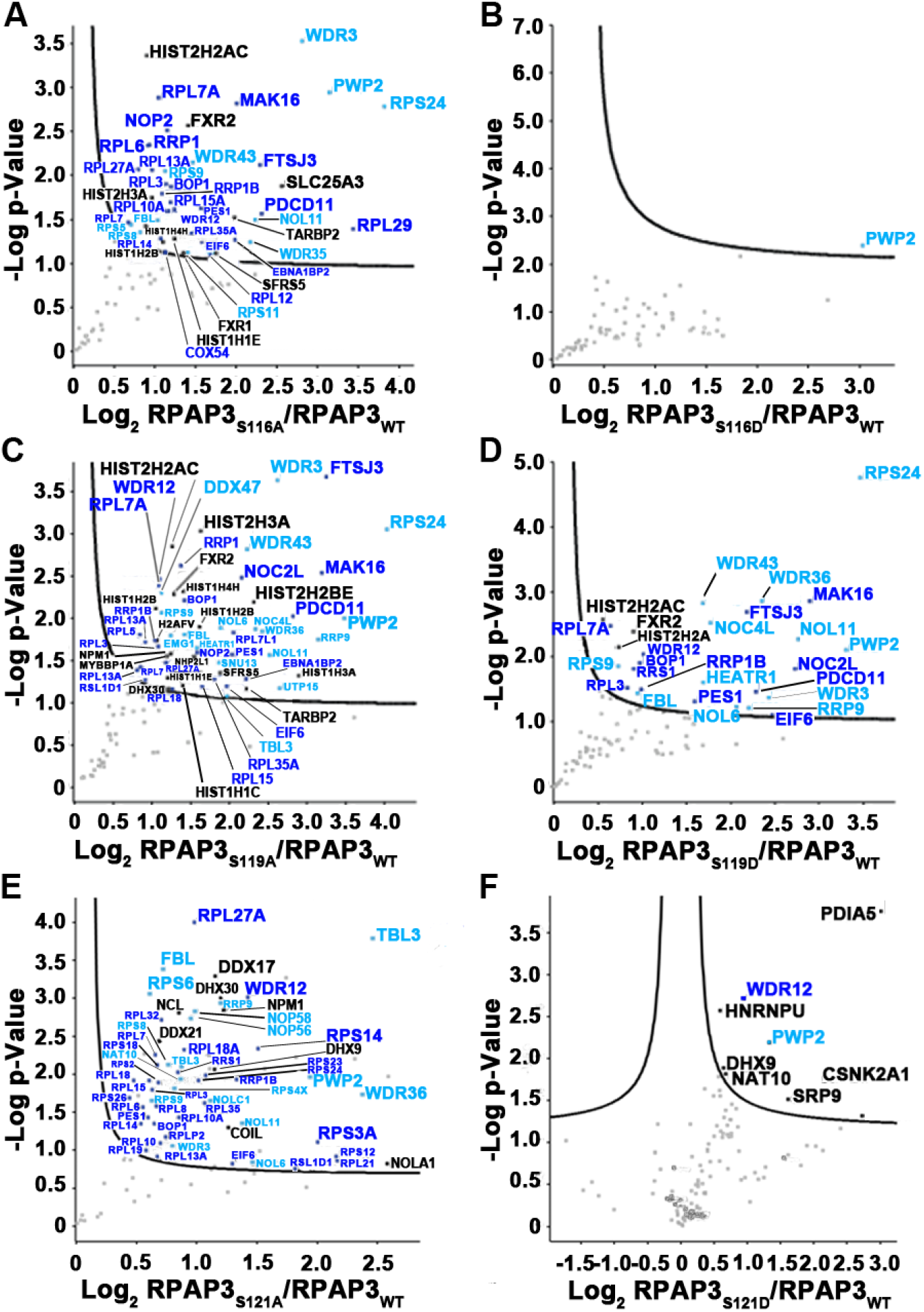
Phospho-null mutation of RPAP3 phosphosites S116, S119 and 121 increase its association with preribosomal proteins. FLAG-tagged RPAP3-WT or the phospho-null (A) RPAP3-S116A, (C) RPAP3-S119A, (E) RPAP3-S121A, or phosphomimetic (B) RPAP3-S116D, (D) RPAP3-S119D, or (F) RPAP3-S121D were expressed in HEK293T cells for 48 h, purified with an anti-FLAG antibody and digested with trypsin. Purifications were performed in triplicate and the co-purified proteins were identified by liquid chromatography-tandem mass spectrometry (LC-MS/MS). The label-free quantification (LFQ) intensity of each peptide was computed via MaxQuant (Version 1.6.2.10) against the charactarized Uniprot database (updated on June 3th 2018) and further analyzed by Perseus (Version 1.6.1.3). Volcano plots illustrate the log_2_-transformed average LFQ-intensity difference between mutant and WT (x axis), and the –log_2_ p value obtained via a two-tailed t test adjusted with a permutation-based multiple hypothesis testing with 10,000 iterations and an s0 correction factor of 0.1 (y axis). All mutants showed significant association with ribosomal proteins and biogenesis factors associated with small (40S; light blue) and large (60S; dark blue) subunits of the ribosome. All other proteins that cleared False Discovery Rate (FDR) threshold of 0.05 are marked in black.

Figure 3 shows results of an analysis of unique and common interactors of RPAP3 S116 and S119 mutants obtained in our AP-MS experiments, highlighting association of several proteins with Gene Ontology enrichment in the ‘Biological Process’ category of Ribosome Biogenesis, both small (40S) and large (60S) subunits (Fig. 3A, 3B). An interaction network of enriched proteins in all RPAP3 phosphosite mutants is presented in Figure 3C. Notably, no increase in association with ribosome biogenesis factors was observed when AP-MS experiments were performed with phospho-null or phosphomimetic mutation at S87 of RPAP3 (Supplementary Fig. S1 and Supplementary Table S5). Together, these results indicate that unphosphorylated forms of RPAP3 at S116, S119 and S121 preferentially associate with preribosomal complexes.

**Fig 3.**
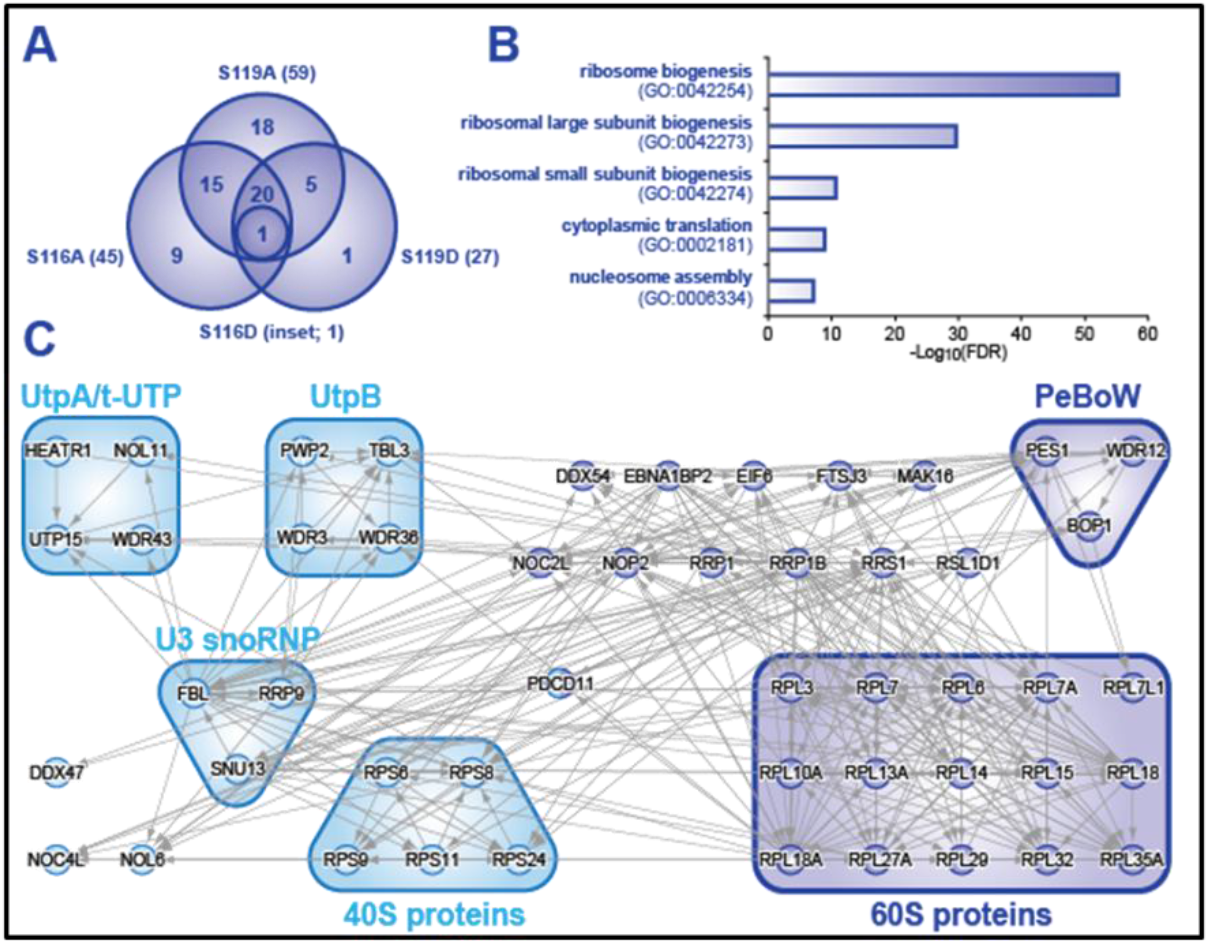
Analysis of interactors of FLAG-tagged RPAP3 phosphosite mutants shows significant enrichment of ribosomal proteins and ribogenesis factors. A) Venn diagram illustrating the number of unique and common interactors identified by AP-MS for RPAP3 S116 and S119 mutants. B) Gene Ontology enrichment analysis in ‘Biological Process’ category for enriched proteins in all RPAP3 phosphosite mutants. The Benjamini-Hochberg FDR was computed using the PANTHER (version 14.1) statistical overrepresentation test. C) Interaction network of enriched proteins in all RPAP3 phosphosite mutants. The dataset was generated using the Agile Protein Interaction DataAnalyzer (APID) and the graphical output of the network was built using Cytoscape (version 3.7.1). Outliers that did not connect with the rest of the networks were removed. Ribosomal proteins and biogenesis factors associated with small (40S) and large (60S) subunits of the ribosome are shown in light and dark blue, respectively.

### CK2 phosphorylates RPAP3 at Ser116, Ser119 and Ser 121

The presence of CK2 recognition sites surrounding RPAP3 S116, 119 and 121 suggests that CK2 is the kinase responsible for phosphorylation of these residues. If this is indeed the case, we would expect that CK2 inhibitors, by favoring an unphosphorylated state, would also favor recruitment of preribosomal assembly factors. To test this hypothesis, we performed AP-MS experiments using wildtype Flag-tagged RPAP3 expressed in cells treated with the CK2 inhibitors CX-4945 or TBB as compared to vehicle only (DMSO). Independent treatment with each inhibitor increased the association of wildtype RPAP3 with ribosomal biogenesis proteins (Fig. 4; Supplementary table S6). CX-4945- and TBB-treated cells increase RPAP3 association with ribosomal proteins and biogenesis factors associated with small (40S; light blue) and large (60S; dark blue) subunits of the ribosome. These results indicate that maintenance of RPAP3 in the unphosphorylated state through the use of CK2 inhibitors increases the association with ribosomal biogenesis factors.

**Fig 4.**
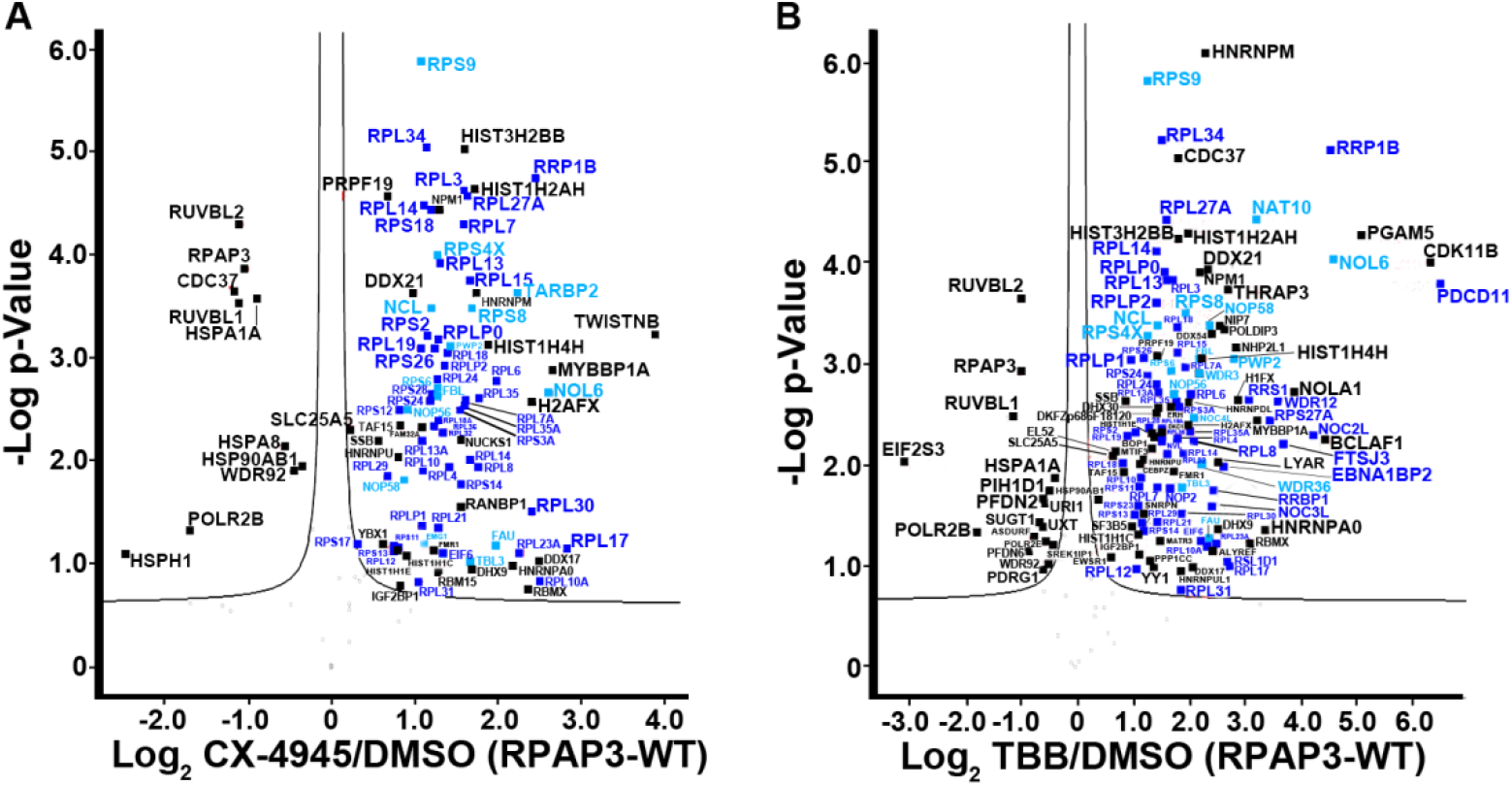
Treatment with two different CK2 inhibitors, CX-4945 and TBB, increases the association of wild-type RPAP3 with ribosomal biogenesis protein. FLAG-tagged RPAP3-WT were expressed in HEK293T cells for 48 h and treated overnight with either (A) 10µM of CX-4945 or (B) 100µM of TBB. The drug-treated samples were purified with an anti-FLAG antibody, and digested with trypsin. Purifications were performed in triplicate and the co-purified proteins were identified by liquid chromatography-tandem mass spectrometry (LC-MS/MS). The label-free quantification (LFQ) intensity of each peptide was computed via MaxQuant (Version 1.6.2.10) against the characterized Uniprot database (updated on June 3th 2018) and further analyzed with Perseus (Version 1.6.1.3). Volcano plots illustrate the log_2_-transformed average LFQ-intensity difference between cells treated with the inhibitors as compared to vehicle only (DMSO) (x axis), and the –log_10_ p value obtained via a two-tailed t-test adjusted with a permutation-based multiple hypothesis testing with 10,000 iterations and an s0 correction factor of 0.1 (y axis). Proteins marked in black, dark blue and light blue are considered significantly different.

To confirm that CK2 is indeed the kinase responsible for RPAP3 S116, S119 and S121 phosphorylation, we purified each phospho-null RPAP3 mutant individually, the triple AAA mutant and wildtype RPAP3 using GST pulldown from E. coli and used the GST-cleaved purified recombinant proteins in CK2 phosphorylation assays *in vitro*. The data indicates that only the triple mutant is resistant to phosphorylation, suggesting that CK2 is indeed responsible for RPAP3 phosphorylation, a process that is not inhibited through substitution of a single residue at a time (Fig. 5).

**Fig 5.**
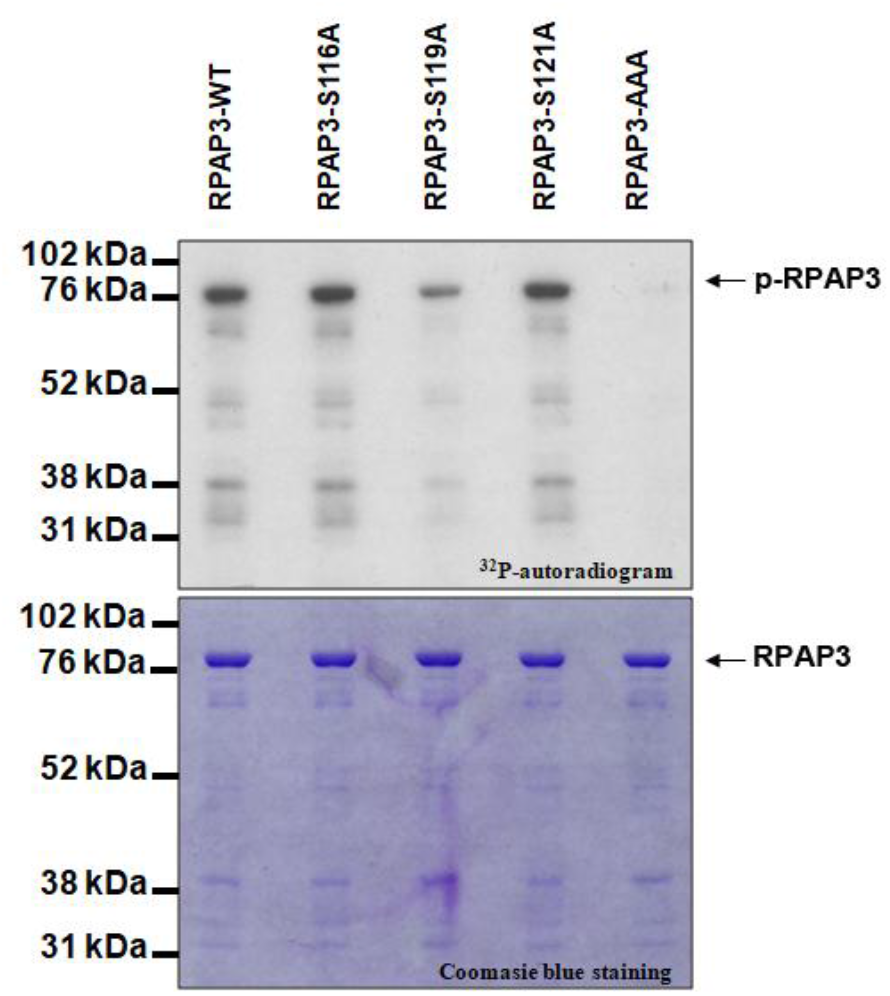
Alanine substitution of RPAP3 phosphosites collectively hinder its phosphorylation by CK2 *In Vitro*. RPAP3 WT or alanine substitution (S116A, S119A, S121A and AAA triple mutant) were expressed in BL-21 and purified by GST-pulldown. 2µg of GST-Cleaved purified RPAP3s were incubate in presence of recombinant CK2 and ^32^P-radiolabelled ATP and loaded on SDS-PAGE after reaction. Autoradiography is shown on the top and the coomasie blue staining control is shown on the bottom panel.

### Assembly of preribosomal complexes is impaired by PAQosome subunit silencing

To confirm a role of the PAQosome in preribosomal complex assembly, we proceeded to the single silencing of two PAQosome subunits, RPAP3 or WDR92, and the monitoring of its effect on the interactomes of the ribosomal biogenesis factors RRP1B and PDCD1. Of note, RPAP3 is part of the PAQosome core module and WDR92 is a component of the prefoldin-like (PFDL) module. The results indicate that silencing of RPAP3 or WDR92 both increase the association of ribosome biogenesis factors RRP1B and PDCD11 with cytokine MDK^26^. However, a decreased interaction with helicase DDX21^25^ was only observed when WDR92 was silenced (Fig. 6; Supplementary Tables S7 and S8).

**Fig 6.**
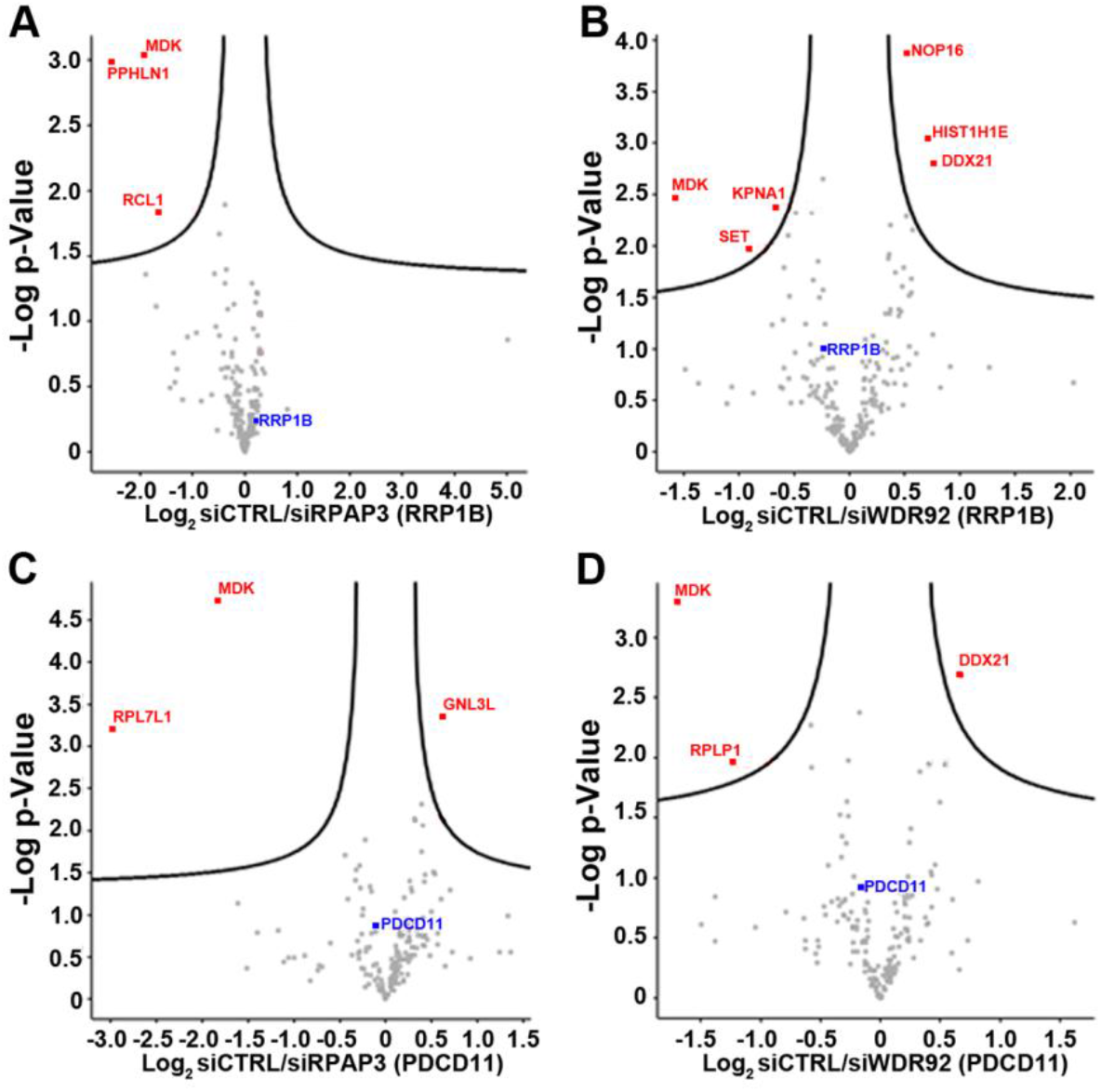
Knockdown of PAQosome subunits RPAP3 and WDR92 increases the association of ribosome biogenesis factors RRP1B and PDCD11 with cytokine MDK (and decreases interaction with helicase DDX21 in the case of WDR92 KD). FLAG-tagged-RRP1B and - PDCD11 were expressed in HEK293T cells and treated a siRNA against RPAP3 (A, C) or WDR92 (B, D). The samples were purified with an anti-FLAG antibody and digested with trypsin. Purifications were performed in triplicate and the co-purified proteins were identified by liquid chromatography-tandem mass spectrometry (LC-MS/MS). The label-free quantification (LFQ) intensity of each peptide was computed via MaxQuant (Version 1.6.2.10) against the charactarized Uniprot database (updated on June 3th 2018) and further analyzed with Perseus (Version 1.6.1.3). Volcano plots illustrate the log_2_-transformed average LFQ-intensity difference between siControl over targeted siRNA-treated cells (x axis), and the – log_10_ p value obtained via a two-tailed t-test adjusted with a permutation-based multiple hypothesis testing with 10,000 iterations and an s0 correction factor of 0.1 (y axis). The bait is marked in blue and the proteins marked in red are considered significantly different between the conditions.

Moreover, siRNA against PDRG1, URI1 and UXT of the prefoldin-like complex of the PAQosome affect the proper assembly of the preribosome (Fig S2, Supplementary Table S9). These results indicate that perturbations in the abundance of PAQosome subunits interfere with association of ribosome biogenesis factors, strengthening the conclusion that the PAQosome potentially regulates ribosome biogenesis.

## Discussion

In this manuscript, we provide evidence that phosphorylation of the core PAQosome subunit RPAP3 modulates binding of several cellular factors, mainly the ribosome subunits and their assembly factors. This finding describes a third mechanism of substrate selection by this HSP90 co-chaperone. With the two other mechanisms, namely the use of specific adaptors^4–6, 9, 11, 13, 27^ and the integration of alternative subunits^16^, posttranslational modification of subunits has the ability to multiply by many fold, if used in combination with the other mechanisms, the complexity of substrates for the PAQosome. Based on this knowledge, future studies will likely identify several additional clients for this complex-assembly nanomachine.

Biogenesis of the ribosome is a highly complex process where assembly factors are added at specific stages and then removed to favor entry of new assembly factors as reviewed in yeast^28^. This process requires approximately 200 ribosomal proteins and structural rRNAs^29^. This is true for both 40S and 60S subunits and takes place mainly in the cell’s nucleolus. In the current study, we show that the PAQosome subunit RPAP3 possesses several serine residues within CK2 consensus site that can be phosphorylated *in vitro* by CK2. These residues are found in proximity of a sequence with nucleolar localization signal (NoLS) homology^30^. These phosphorylation sites modulate association of multiple ribosomal assembly factors with ribosome subunits, suggesting a complex role of the PAQosome in ribosome assembly, possibly by acting at various stages of ribosome biogenesis. It is interesting to note that a single kinase, CK2, phosphorylates all three RPAP3 Ser residues involved in preribosomal complex assembly. Pharmacological inhibition of this kinase, by favoring the unphosphorylated status of RPAP3, increased association with preribosomal complexes. Whether other RPAP3 (or other PAQosome subunits) residues are involved in dictating substrate specificity through phosphorylation/dephosphorylation involving CK2 (or any other kinase) is not known at this stage. This is supported by the fact that CK2 was shown to accumulate in the nucleolus of cells^31^ and its kinase activity was already shown to modulate the function of various nucleolar proteins associated with ribosomal biogenesis^32, 33^, CK2 can phosphorylate RNA polymerase I and block its transcriptional activity thus, altering ribosome biogenesis^34^. Moreover, CK2 can also stimulate rRNA expression after AKT signaling pathway activation^35^. RPAP3 phosphorylation could be another layer of regulation to tightly control ribosomal biogenesis^36^.

On the hand, accumulation of MDK^26^, an essential factor shown in 45S precussor expression after RPAP3 or WDR92 inhibition is another clue that supports the role of the PAQosome in ribosomal maturation. Likewise, DDX21 was shown to be a mammalian component of the preribosome required in the late step of 40S ribosome maturation^37^. Downregulation of DDX21 does not block 90S to 40S maturation but affects the loading of two specific snoRNPs in the 40S ribosome^37^. These results support the role of the PAQosome in ribosome assembly. Moreover, the several ribosomal factors observed after silencing of the prefoldin-like complex subunits strengthen the link between the PAQosome and ribosomal maturation. However, not being able to pull-down the PAQosome subunits during AP-MS experiment suggest that the co-chaperone might act as a transient reaction during ribosomal maturation and a cause of AP-MS limitation^38^. Additional studies will be required to decipher the exact function of RPAP3 and other PAQosome subunits along this complex biogenesis process.

## Conclusions

Our results uncovered phosphorylation of the core PAQosome subunit RPAP3 as a third mechanism of client specificity/selectivity by the HSP90 co-chaperone PAQosome. Unphosphorylated RPAP3 at Ser 116, 119 and 121 associates with several ribosomal assembly factors and ribosome subunits. These phosphorylation events are catalyzed by kinase CK2. Taken in the context of previous reports, our data support the notion that the PAQosome has evolved with multiple distinct mechanisms to control client/substrate specificity in humans.

## Acknowledgments

This work was supported by a grant from the Ministry of Economy and Innovation form the Government of Quebec. B.C. holds the IRCM Bell-Bombardier Research Chair. We are grateful to Denis Faubert, Josée Champagne, Sylvain Tessier and Marguerite Boulos for their assistance in sample processing and mass spectrometric data analysis.

## Notes

### Competing Interest Statement

The authors have declared no competing interest.

