## Supplementary information. for "The unphosphorylated form of the PAQosome core subunit RPAP3 binds ribosomal preassembly complexes to modulate ribosome biogenesis"

### Supplementary Figures

**Fig S1 – Phospho-null mutation of RPAP3 phosphosite S87 does not increase its association with preribosomal proteins.**
**Fig S2 – Silencing of PAQosome subunits PDRG1, URI1, and UXT alter the ribosome biogenesis factor RRP1B's interactome.**

### Supplementary Tables

**Table S1.** Plasmids and oligonucleotides used in this study.
**Table S2.** Filtered protein list from MaxQuant label-free quantification analysis of RPAP3-S116A and S116D experiment related to Fig 2A and 2B.
**Table S3.** Filtered protein list from MaxQuant label-free quantification analysis of RPAP3-S119A and S119D experiment related to Fig 2C and 2D.
**Table S4.** Filtered protein list from MaxQuant label-free quantification analysis of RPAP3-S121A and S121D experiment related to Fig 2E and 2F.
**Table S5.** Filtered protein list from MaxQuant label-free quantification analysis of RPAP3-S87A and S87D experiment related to Fig S2A and S2B.
**Table S6.** Filtered protein list from MaxQuant label-free quantification analysis of DMSO, CX4945 and TBB treated cells related to Fig 4A and 4B.
**Table S7.** Filtered protein list from MaxQuant label-free quantification analysis of 3X-FLAG-RRP1B interactome in siRNA against RPAP3 and WDR92 treated cells related to Fig 6A and 6B.
**Table S8.** Filtered protein list from MaxQuant label-free quantification analysis of 3X-FLAG-PDCD11 interactome in siRNA against RPAP3 and WDR92 treated cells related to Fig 6C and 6D.
**Table S9.** Filtered protein list from MaxQuant label-free quantification analysis of 3X-FLAG-RRP1B in siRNA against UXT, URI1, and PDRG1 treated cells related to Fig S3A, S3B and S3C.

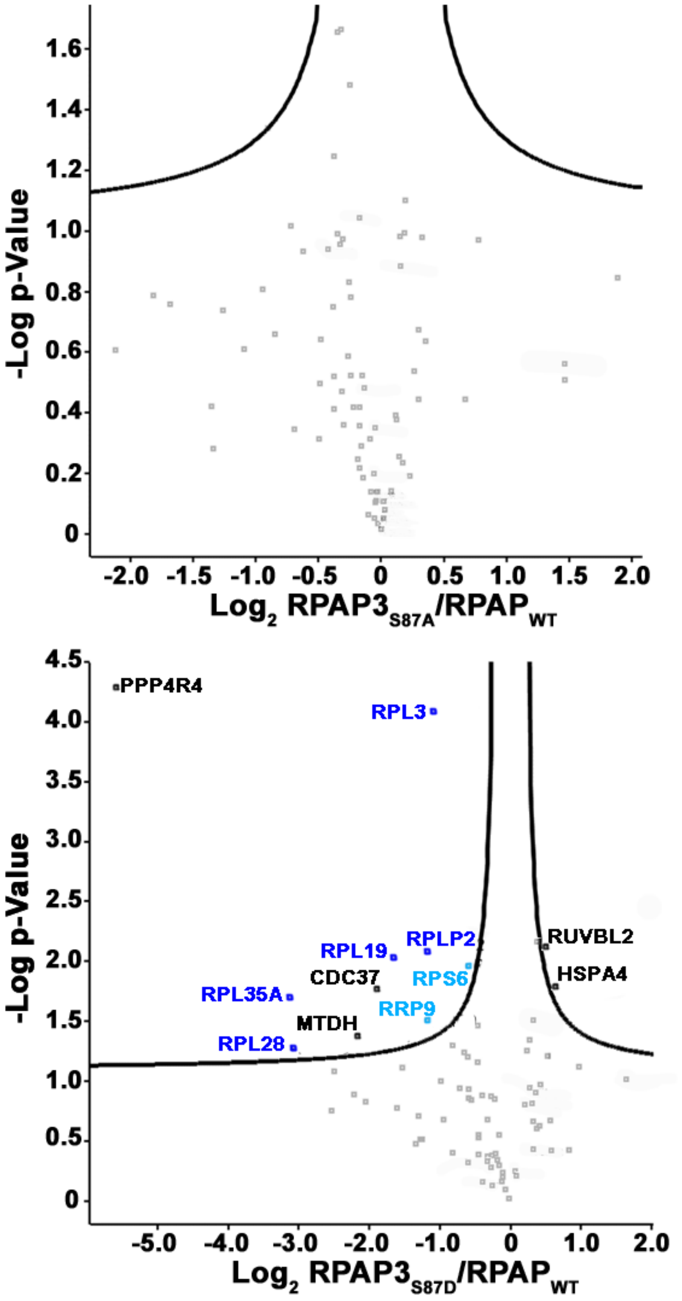


**Fig S1 - Phospho-null mutation of RPAP3 phosphosite S87 does not increase its association with preribosomal proteins.**

FLAG-tagged RPAP3-WT and (A) FLAG-tagged RPAP3-S87A or (B) FLAG-tagged RPAP3-S87D were expressed in HEK293T cells for 48 h, purified with an anti-FLAG antibody, and digested with trypsin. Purifications were performed in triplicate and the co-purified proteins were identified by liquid chromatography-tandem mass spectrometry (LC-MS/MS). The label-free quantification (LFQ) intensity of each peptide was computed via MaxQuant (Version 1.6.2.10) against the charactarized Uniprot database (updated on June 3th 2018) and further analyzed by Perseus (Version 1.6.1.3). Volcano plot illustrates the $log_2$-transformed average LFQ-intensity difference between mutant and WT (x axis), and the $–log_{10}$ p value obtained via a two-tailed t test adjusted with a permutation-based multiple hypothesis testing with 10,000 iterations and an s0 correction factor of 0.1 (y axis). Identified proteins were significantly different with a False Discovery Rate (FDR) threshold of 0.05. Ribosomal proteins and biogenesis factors associated with small 40S and large 60S subunits of the ribosome were marked in light blue and dark blue respectively. Other significantly different proteins were marked in black.

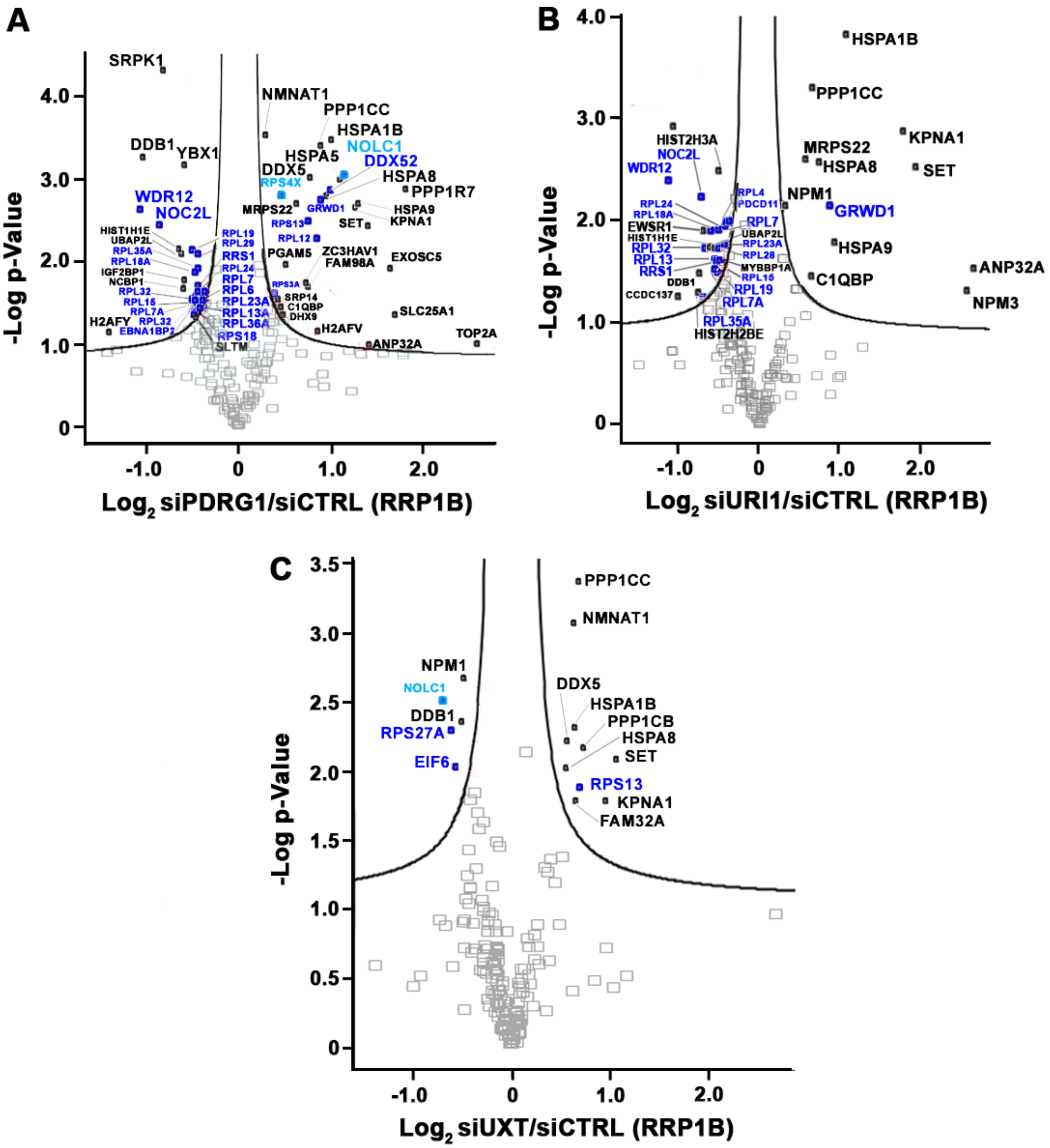


**Fig S2 – Silencing of PAQosome subunits PDRG1, URI1, and UXT alter the ribosome biogenesis factor RRP1B's interactome.**

FLAG-tagged RRP1B was expressed in HEK293T cells treated with a siRNA against (A) PDGR1, (B) URI1 or (C) UXT. The FLAG-tagged samples were purified with an anti-FLAG antibody, and digested with trypsin. Purifications were performed in triplicate and the co-purified proteins were identified by liquid chromatography-tandem mass spectrometry (LC-MS/MS). The label-free quantification (LFQ) intensity of each peptide was computed via MaxQuant (Version 1.6.2.10) against the charactarized Uniprot database (updated on June 3th 2018) and further analyzed by Perseus (Version 1.6.10.43). Volcano plots illustrate the $\log_2$-transformed average LFQ-intensity difference between mutant and WT (x axis), and the –$\log_{10}$ p value obtained via a two-tailed t test adjusted with a permutation-based multiple hypothesis testing with 10,000 iterations and an s0 correction factor of 0.1 (y axis). Identified proteins were significantly different with a False Discovery Rate (FDR) threshold of 0.05. Ribosomal proteins and biogenesis factors associated with small 40S and large 60S subunits of the ribosome were marked in light blue and dark blue respectively. Other significantly different proteins were marked in black.
